# Perceiving surface reflectance requires attention

**DOI:** 10.1101/2025.10.14.682481

**Authors:** Erin Goddard, Kavita Paul Remician

## Abstract

Colour constancy refers to our ability to distinguish changes in surface properties from changes in the properties of light illuminating the scene. The apparent ease with which we can tell that objects do not change in their surface properties, for example, when moving from sunlight to the shade, belies the complexity of solving this ill-constrained problem. Although there is a substantial body of work testing which image cues might be used to accomplish this, there is surprisingly little known of how the brain performs this computation. Here, we tested a fundamental aspect of the perceptual scission of surface reflectance from illuminant/filter properties: whether it requires attention. We measured visual search times for both surface reflectance (requiring separation of surface and illuminant properties) and local tristimulus value (which does not). We found a clear difference between the two: visual search for colour defined by tristimulus value was fast and near-parallel, while search for a particular surface reflectance was slow and consistent with the serial deployment of attention. That is, search times suggest that the perceptual separation of surface and illuminant or filter properties in colour constancy may require an attention-based process analogous to perceiving conjunctions of simple features in ‘feature binding’. These results offer important new insights into the sequence of processes the brain uses to accomplish colour constancy.

## Introduction

Colour provides a valuable visual cue to the surface properties of objects, which is used in object memory and recognition, and for communication (e.g. Witzel & Gegenfurtner, 2018). However, the spectral composition of light wavelengths that reach the eye depend not only on surface reflectance properties, but on the spectral properties of the light illuminating the scene, and the transmittance properties of any filters through which the surface is viewed. Recognising surfaces across different illumination and/or filter conditions, or ‘colour constancy’, requires the perceptual separation or scission of surface reflectance properties from the properties of the illuminant and any filters. Although colour constancy is integral to understanding human colour vision, and how we use colour information, the neural mechanisms underlying this process remain poorly characterised. Previous work has produced a detailed account of the image cues that affect colour constancy which suggests the kinds of neural computations that the visual system may use, and has identified likely candidates for where in the brain there exist responses to colour that are shifted towards colour constant representations. Here we seek to understand these neural computations from a different angle, using search slopes in a visual search task to measure whether perceived surface reflectance is available preattentively, consistent with unlimited-capacity parallel processing, or if perceiving surface reflectance requires the direction of focal attention.

The computational problem of colour constancy is underconstrained (Smithson, 2005), and our degree of colour constancy is typically imperfect, but varies considerably across scenes (Hansen et al., 2007; Kraft & Brainard, 1999; McCann et al., 1976). Both successes and failures of colour constancy help to constrain our understanding of how the human visual system arrives at a ‘best guess’ of surface properties (Brainard et al., 2006; Gilchrist et al., 1999). A range of visual cues have been shown to improve constancy, implying that they are used by the visual system to perceptually separate surface and illuminant properties. These include lower-level cues that might be computed from relatively simple image properties, without first inferring the 3D structure, such as the ratio of intensity values in different lightness channels across the scene (Land & McCann, 1971) and the colour of the brightest point in the scene (Gilchrist et al., 1999). However, such global scene statistics are not sufficient to fully account for colour constancy (Kraft & Brainard, 1999; Smithson, 2005), and a number of higher-level cues have also been shown to influence colour constancy, such as the presence of interreflections (Bloj et al., 1999), specular highlights (Anderson & Kim, 2009; D’Zmura & Lennie, 1986) and memory of object colours (Granzier & Gegenfurtner, 2012). Given that colour constancy improves with cues that depend upon extraction of the scene’s 3D structure, or object recognition, it seems likely that perception of surface reflectance may be a relatively high-level visual feature, emerging at a later stage of visual processing.

There is psychophysical (Goddard et al., 2010), electrophysiological (Walsh et al., 1993; Wild et al., 1985; Zeki, 1983) and neuroimaging (Albers et al., 2022; Bannert & Bartels, 2017; Bartels & Zeki, 2000; Zeki & Marini, 1998) evidence of cortical responses that are consistent with a shift towards surface reflectance, and away from the local tristimulus coordinates of light reaching the retina. While these results demonstrate that the brain represents surface reflectance, and suggest where this may occur (e.g. area V4), this does not constrain the neural processes by which these representations emerge.

Specifically, these representations of surface reflectance properties may occur automatically and in parallel across the visual field; alternatively, the process of separating surface and illuminant properties may rely on processes that are contingent upon attentional selection, in a manner qualitatively similar to the binding of some feature conjunctions, which require limited-capacity, attention-driven mechanisms (Treisman, 1996; Treisman & Gelade, 1980; Wolfe & Cave, 1999).

In this study, we tested whether the perceptual scission of surface and filter properties (colour constancy) requires the engagement of limited-capacity, attention-driven mechanisms. To do this, we compared the effectiveness of surface reflectance and local tristimulus values in their ability to guide attention in a visual search task. Visual search tasks have an extensive history of being used to separate preattentive features from those requiring attention (Wolfe, 2021). In general terms, preattentive features can be detected rapidly via an almost parallel process across the entire visual field, whereas many feature conjunctions or complex features require some degree of slower, serial search, taking longer with increasing set size. Treisman and Gelade’s (1980) seminal ‘Feature Integration Theory’ posited that targets defined by simple features (e.g. colour or shape of a letter) could be detected similarly quickly regardless of the set size since colour and shape are both registered automatically and in parallel across the entire visual field, whereas finding a conjunction of features (e.g. a blue letter ‘S’) requires attention to be directed serially to different locations, acting as a ‘glue’ that binds the otherwise free-floating feature signals into a unified object percept. There is not a strict dichotomy between preattentive, completely parallel processing and item-by-item serial search: these form two ends of a continuum of ‘search efficiency’, rather than discrete categories (e.g. Lleras et al., 2020; Thornton & Gilden, 2007; Wolfe, 2021). Wolfe’s (1994, 2021) ‘Guided Search’ models formalise how the search efficiency of the target feature dimension interacts with a range of factors, including other feature dimensions, and top-down influences, to produce a continually updating priority map, which predicts how attention will be guided during different visual searches. Of central importance to the present study, how efficiently a feature dimension guides attention during visual search is not simply a measure of how slow or difficult it may be to perceive the relevant feature (as captured by a measure such as reaction time when considering a single object); instead, the efficiency of visual search also gives insight into whether perception of the feature requires a capacity-limited process which produces a tight selection bottleneck, such as feature binding, where only one or very few items that are current objects of attention can be processed simultaneously (Wolfe, 2021). Here we used visual search to directly compare two feature dimensions, surface reflectance and tristimulus coordinates, in their capacity to guide search efficiently, using search tasks that were as closely matched as possible for the different dimensions.

Colour is an effective simple feature for efficient visual search and can be used as a fast and preattentive guide to search when there is sufficient colour difference between a target and the distractor items (e.g. Carter, 1982; Duncan & Humphreys, 1989; Nagy & Sanchez, 1990). However, these studies use elements where perceived surface properties and tristimulus coordinates are not dissociated, meaning it is unclear whether either or both of local tristimulus values and surface reflectance properties can guide search preattentively. Local tristimulus coordinates, or the spectral properties of light reaching the eyes, are of less behavioural relevance than perceived surface and illuminant properties, and may not be represented once scission has occurred. This possibility is demonstrated in illusory percepts that can be accounted for by scission (e.g. Anderson et al., 2011), and is illustrated in Figure 1**A**. In example scenes *iii* and *iv* (Figure 1**A**), the dragon objects are matched in their tristimulus coordinates but are typically perceived as differently coloured. If asked to judge tristimulus values, observers may rely on explicit reasoning rather than perceptual experience. This illustrates the importance of considering how a participant interprets their task when investigating the perception of stimuli that can be perceived as a surface under a translucent filter or coloured illuminant: whenever the participant perceives multiple superimposed ‘layers’ of information (such as those corresponding to surface reflectance, filter transmittance and illuminant properties), there are multiple approaches for reporting their percept (e.g. Ekroll et al., 2004). For instance, when instructions are explicitly varied to encourage participants to make matches based on surface properties or tristimulus coordinates, participants can adjust their responses appropriately (Arend & Reeves, 1986; Cornelissen & Brenner, 1995; Troost & de Weert, 1991). However, a further complication is the complexity of separating perception from inference in such tasks. Radonjic & Brainard (2016) systematically varied instructions across participants, and found lower individual variability when participants matched tristimulus values rather than surface reflectance properties. They inferred that, for their stimuli, the tristimulus matches may have better reflected perceptual experience, while judging surface reflectance likely included an adjustment based on cognitive inference. They note that a similar effect could potentially occur in the opposite direction: for stimuli where colour constancy is high, instructions asking for a tristimulus match could prompt participants to reason that they must adjust their match in the direction opposite of constancy (as taught in courses on topics such as painting). Hence if scission occurs rapidly and preattentively, perceived surface reflectance would be available preattentively, but searching for a specific tristimulus value may require a slower, limited-capacity reasoning process to ‘recombine’ the surface and illuminant or filter information and infer the local tristimulus value. Alternatively, if scission requires a limited-capacity attentional mechanism, then searching for a specific surface reflectance should be slower and serial. In this scenario, before attention is deployed and scission has occurred, the colour information available preattentively should be closer to the tristimulus coordinates, meaning that tristimulus value search would be closer to parallel. To distinguish between these alternatives, we measured visual search times for both surface reflectance search and tristimulus value search, using stimuli where these features were dissociated. We found patterns of reaction times consistent with tristimulus values being available preattentively, but the perception of surface reflectance properties requiring the serial deployment of attention.

**Figure 1:**
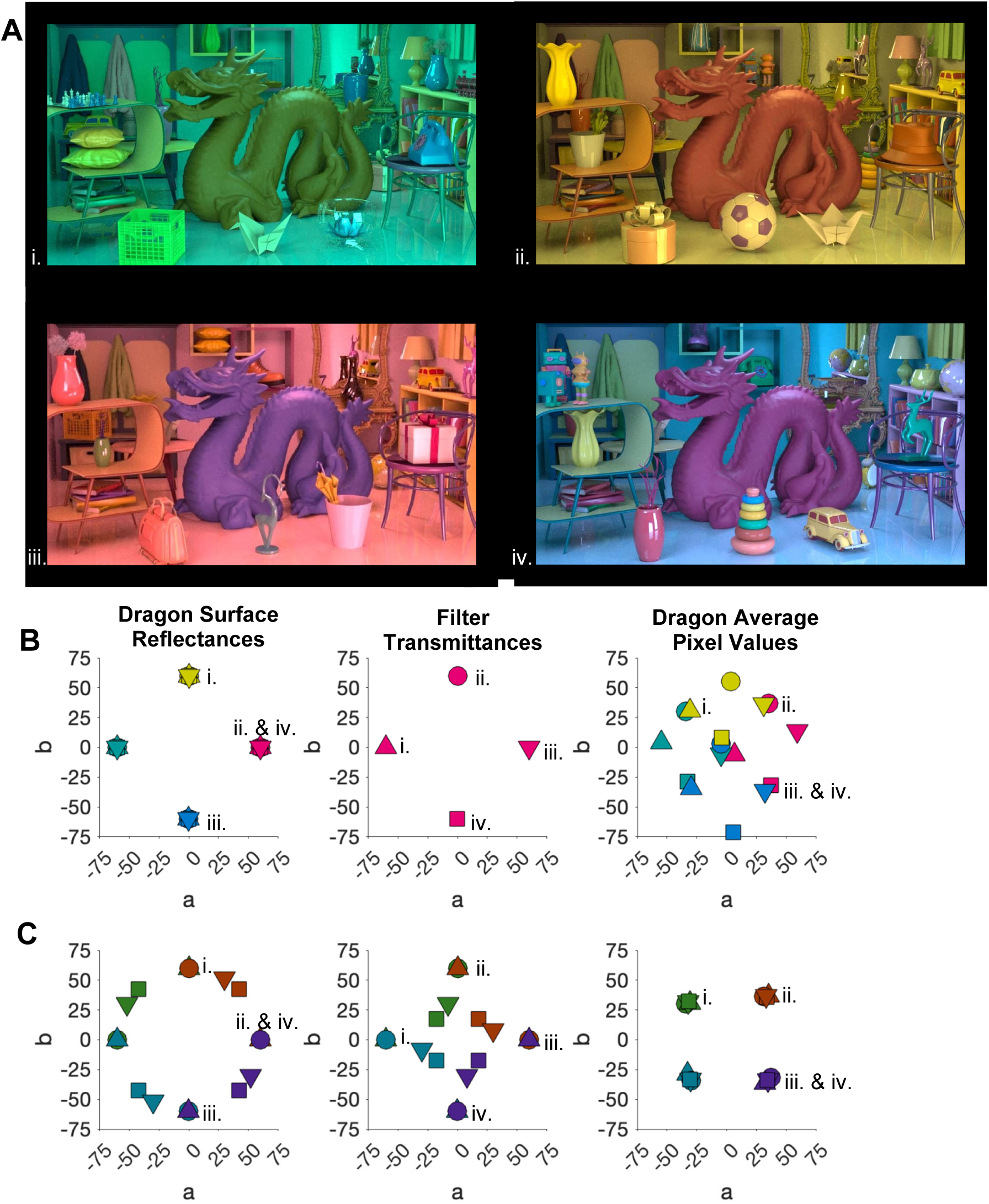
Example stimuli. **A**: Four example scenes used as stimuli, simulating 4 filters, and central dragon objects of 3 different surface reflectances. **B&C**: The way in which surface reflectance properties and local pixel values of the central dragon were dissociated here is illustrated in the CIE L*a*b* coordinates of stimuli used in the discrimination and visual search tasks. Stimuli had defined surface reflectances (**B&C**, left) and filter transmittances (**B&C**, middle) which yielded average pixel values (**B&C**, right) in the rendered stimuli that were approximately midway between the surface and filter. **B** depicts 16 stimuli of 4 unique surface reflectances (used in the surface reflectance versions of the discrimination and search tasks), and **C** shows 16 stimuli of 4 unique tristimulus coordinates (used in the tristimulus versions of these tasks). 8 of the 16 stimuli, including the 4 in **A**, are common to both **B** and **C**. In **B** the markers are coloured according to the dragon’s surface reflectance in each stimulus, and marker shape varies with filter transmittance. In **C** markers are coloured according to the dragon’s tristimulus values; for each set of tristimulus coordinates there were 4 different combinations of dragon surface and filter properties, shown in 4 different marker shapes. Both **B** and **C** are annotated to indicate the markers corresponding to the example scenes in **A**.

## Materials and Methods

### Overview

We conducted two experiments using physically realistic, complex stimuli designed to dissociate surface reflectance properties from local pixel tristimulus coordinates as much as possible (as shown in **Figure 1**). In Experiment 1, we measured participants’ discrimination accuracy for changes in surface reflectance, and changes in tristimulus coordinates, of the central dragon object. By completing this task first, we ensured that all participants were familiar with the stimuli and understood the concepts of surface reflectance and local pixel tristimulus coordinates before proceeding to Experiment 2 (visual search tasks). In Experiment 1, we manipulated stimulus complexity, measuring discrimination sensitivity for both tasks for the original stimuli along with simplified two-tone versions (‘reduced’ stimuli), and, as a control condition, a ‘single colour’ stimulus (tristimulus task only), as shown in **Figure 2**. Discrimination accuracies for the complex stimuli showed that participants were very close to ceiling performance in both tasks when the dragon singletons were separated from the distractors in L*a*b* space by the differences shown in Figure 1 (i.e. surface reflectance differences of those shown in Figure 1**B**, left panel, or tristimulus coordinate differences of those in Figure 1**C**, right panel). In Experiment 2, the same participants performed a visual search for a central dragon of either a specific surface reflectance or a specific tristimulus value. For each search task, all elements were complex scenes and the dragons had one of the 4 unique values in Figure 1 (the 4 values in the left panel of Figure 1**B** for the surface reflectance search or the 4 values in the right panel of Figure 1**C** for tristimulus search), to ensure that the stimulus differences in the relevant dimension were well above threshold, and approximately matched across tasks. We compared the visual search results against three models based on Guided Search 6.0 (GS6) (Wolfe, 2021), where the models assumed that either surface reflectance (but not tristimulus value) guided search preattentively, or vice versa, or that only the tristimulus value of the filter guided search preattentively.

**Figure 2:**
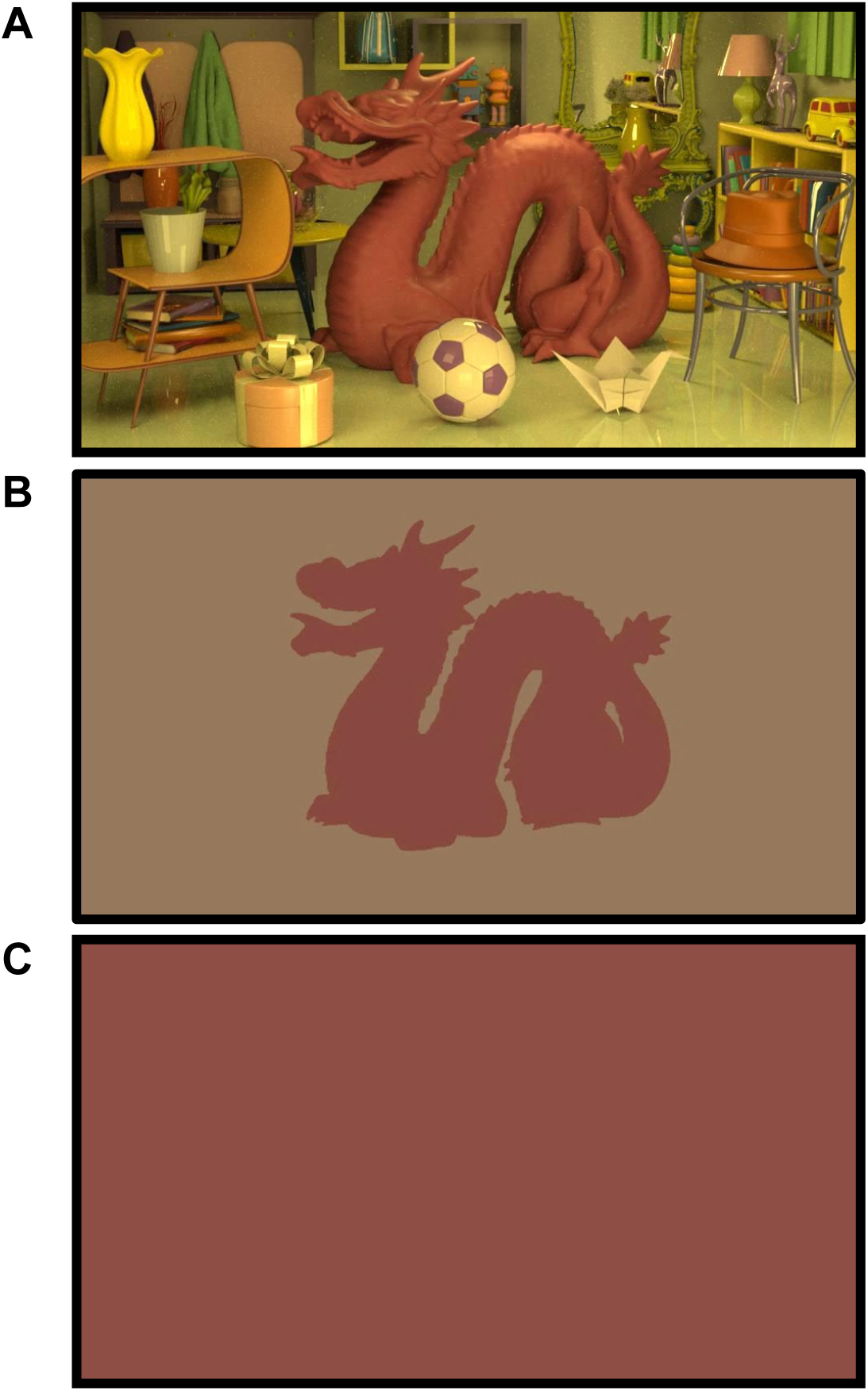
Stimulus conditions in discrimination tasks. **A**: Complex stimulus. **B**: Reduced stimulus. **C**: Single colour stimulus. For each complex stimulus, we created an equivalent reduced and single-colour stimulus. The reduced stimuli were generated by averaging all the XYZ values of all dragon pixels in the corresponding complex stimulus, and using this average for the dragon silhouette. The background of the reduced stimulus was the average of all XYZ values of non-dragon pixels in the complex stimulus. The single colour stimulus was a uniform patch of the same value as the dragon silhouette in the reduced stimulus.

### Stimulus Generation

In designing our stimuli, we aimed to create complex scenes that were rich in cues that could improve colour constancy. All complex stimuli were simulated scenes including a central dragon object in a room with several other objects and items of furniture. Example complex stimuli are shown in **Figure 1** and **Figure 2A**. Four rectangular emitters (light sources) illuminated the scene from the front and above. No emitters were directly visible, but reflections of the emitters were visible on glossy objects in some scenes. Although we originally planned to systematically manipulate illuminant spectral power distributions across stimuli, we found that in the physically realistic rendering process (described immediately below) the illuminant’s spectral power and the central dragon’s surface reflectance properties interacted to produce the dragon’s tristimulus coordinates in ways that made it impractical to define a stimulus set where the dragon’s surface reflectance properties and tristimulus coordinates were dissociated. Since dissociating surface reflectance from tristimulus coordinates was of crucial importance for the design of our visual search task, we kept the spectral power of the light sources constant across all stimuli (a white light with equal power at all wavelengths) and instead simulated a translucent filter located directly in front of the camera, which had varying spectral transmittance properties. Using these filters, the average pixel value of each dragon was approximately midway between the surface and the filter in its L*a*b* coordinates (described in further detail under ‘Dragon and filter properties’, below, and illustrated in **Figure 1B-C**). Varying filter transmittance is not physically identical to varying the illumination; however, making judgements of surface reflectance properties across varying filters requires a perceptual scission of filter and surface properties, which is conceptually similar to a typical colour constancy task where surface reflectance properties are perceptually separated from the illuminant’s spectral properties. We consider this point further in our general discussion.

All scene objects (each item at each possible location) were defined in the open-source 3D graphics software Blender v2.83.1 (The Blender Foundation, Amsterdam, The Netherlands, blender.org). The central dragon object was sourced from the Stanford University Computer Graphics Laboratory (retrieved from graphics.stanford.edu/data/3Dscanrep/, May, 2020), all remaining objects and furniture items were public domain (CC0) objects (retrieved from blendswap.com, May, 2020). Each stimulus was rendered using Mitsuba 2 (Nimier-David et al., 2019), a physically realistic rendering software that simulates paths of light rays within a scene and how they interact with objects based on surface properties and geometry. All surface reflectance properties and emitter (light source) spectral power distributions were defined across the visible light spectrum from 380-780 nm, in 5 nm steps. Scenes were rendered as multispectral images before being saved as the CIE tristimulus (XYZ) values at each pixel. Each image was 960 x 540 pixels in resolution. For display, these were converted to RGB values according to each device’s colour characterisation.

### Scene layout

For every stimulus, participants were required to make judgements about the colour of the dragon object (about either its surface reflectance properties or its tristimulus value, described below under ‘Design’ for both experiments). All other items in the scene were not directly relevant to the tasks but provided additional cues to the scene lighting and filter conditions. The dragon, the walls and many furniture items remained in constant locations for every stimulus, but there were 15 locations that were filled with one of 29 possible objects, that varied randomly across stimuli. No stimulus included the same object more than once, and objects varied in their rotation at different locations, so that the same object was viewed from different angles across stimuli.

#### Background surface properties

Each unique surface in every object was defined by two factors: its spectral reflectance function, and its surface scattering model, which defines how light interacts with the surface, and corresponds to how shiny or matte the surface appears. The surface scattering model of each surface within each object was constant across all scenes in which the object appeared (e.g. the walls was always matte, the vases were always glossy, and the dragon was always a rough dielectric material, simulating a rough plastic or rubber).

The surface reflectance functions of all surfaces in the scene apart from the central dragon were varied randomly across stimuli, to avoid any one surface becoming reliably diagnostic of the filter condition. An exception to this were the mirror, a small number of black surfaces (e.g. boot sole) and a small number of clear glass surfaces (e.g. glass bowl) that were the same each time they appeared in a stimulus. We deliberately selected objects and furniture items that do not have a single typical colour, so that we could randomly vary surface reflectances without varying scene naturalness, and to reduce any contribution of colour memory.

Surface reflectance functions of all surfaces except the dragon and filter were defined using Munsell colours, derived from spectral measurements of physical Munsell chips (Hiltunen, 1996), shown in Supplementary Figure S1. We divided the complete set of Munsell surfaces into nine groups that according to their *value* (dark: 2.5, 3, or 4; medium: 5, 6, or 7; light: 8, 8.5, or 9) and *chroma* (desaturated: 1, 2, or 4; mid-saturated: 6 or 8; saturated: 10, 12, or 14), and assigned each surface to a group. For example, the walls were set to be light and desaturated, and the soccer ball was composed of both a light, desaturated surface and a dark, saturated surface. Across stimuli, the Munsell surface within the group was randomly selected, so that across stimuli objects, walls and furniture surfaces could be any *hue*, and one of a narrower range of *values* and *chromas*. For each *value*/*chroma* group, most surfaces within that group were selected without replacement, to ensure a diversity of surfaces within every scene. The only exceptions were the books, robots and stacker toy, which contained many surfaces of the same group, and so were assigned surface reflectances that could match others in the same scene.

#### Dragon and filter properties

The surface reflectance of the dragon and the transmittance of the filter were defined using custom spectra so that these properties could be parametrically varied in finer hue gradations than the Munsell colour space would allow. For each custom spectra, we defined target **L*a*b\*** coordinates of the relevant spectra, and converted these to target CIE **XYZ** values. We used spectrally broadband basis functions that isolated **X, Y** and **Z** to obtain spectra of the target value, and confirmed that these exactly matched the target **L*a*b\*** coordinates in each case. The resulting custom spectra are shown in **Supplementary Figure S1**. In total, we defined 40 surface reflectance functions that, when under an equal energy white illuminant, would be distributed around in a circle in the **a*b\*** plane (**L**=55, radius 50) with 40 different hue angles. Of these, 12 surface reflectance spectra (those indicated in Figure 1**B-C**, and shown in Supplementary Figure S1), were used as the ‘base’ surface reflectances in the discrimination tasks (Experiment 1) and in the visual search tasks (Experiment 2). The 12 base surface reflectances included 4 that were used in both surface and tristimulus tasks in Experiments 1-2, and a further 8 surface reflectances that were only used as base surface reflectances for stimuli in the tristimulus tasks (seen in Figure 1B-C, left panels). In the discrimination tasks described below, singletons were either 18, 30, 60, 90 or 180 degrees different from the base surface in hue angle: all combinations of base surface and singleton offset gave 40 unique surface reflectance spectra. We also defined a set of 12 filter transmittance functions with the **a*b\*** values shown in the middle panels of **Figure 1B-C** (full spectra in Supplementary Figure S1), with **L**=85. In a similar way to the surface reflectances, 4 of the filter transmittances were uses in both surface and tristimulus tasks in Experiments 1-2, and the remaining 8 filter transmittances were only used for stimuli the tristimulus tasks. For every combination of dragon surface reflectance and filter transmittance, we created 5 versions with different randomly selected objects and background surfaces: every stimulus we created had a unique randomisation seed.

As mentioned above, although we aimed to create a set of stimuli that where both surface reflectance properties and tristimulus coordinates varied systematically, the tristimulus coordinates of the dragon pixels were not directly controlled. To test how well the a*b* coordinates of the average pixel values were predicted by the average of the a*b* coordinates of the surface and filter used in the render, we measured these empirically, by taking the averages of pixels corresponding to the dragon location, excluding pixels where the dragon was obscured by other objects. Light reflected from the dragon in the rendered scenes depends primarily on the reflectance of the dragon, the spectral power of the light sources, and the transmittance of the filter, but also varies with how light is interreflected amongst other scene objects. For this reason, although the average pixel values of the dragon were approximately equidistant from the surface and filter in a*b* coordinates, there was some variability, as seen in Figure 1**B-C**, where the surface reflectances and filter transmittances are spaced with perfect rotational symmetry (order 4), but the average pixel values do not perfectly conform to this. Across stimuli, Pearson correlations between the predicted and measured a*b* coordinates were very high (r=0.995 for a*, r=0.990 for b*). Any small discrepancies were most relevant to the discrimination tasks (Experiment 1), which we considered in greater detail below (under Design: Experiment 1, Discrimination Tasks).

### Participants

12 participants (9 female, 3 male), aged 18 to 40 (M=25.2 years, SD =7.2 years) took part, including both authors, and 10 participants who were naïve to the purposes of the study. All participants had normal or corrected-to-normal visual acuity and normal colour vision, as assessed by the Hardy-Rand-Rittler pseudoisochromatic plates (4^th^ edition, Richmond Products), which include screening for red-green and tritan colour vision abnormalities. Each participant took part in all experimental conditions, with the total testing time of approximately 4 hours divided across at least 2 sessions, on separate days. One participant withdrew after a single session, after completing only half the conditions, so their data were excluded from all analyses. A second participant did not complete all conditions of the visual search experiment, so their data were only included in the colour discrimination experiment. All participants were compensated for their time at a rate of AU$20 per hour. Each participant provided informed consent, and the entire study was carried out with the approval of the Human Research Ethics Approval Panel C, UNSW (HC3129).

### Stimulus Display

In all experiments, stimuli were displayed on a 32” Display++ LCD Monitor by Cambridge Research Systems (resolution 1920x1080, refresh rate 120 Hz, with integrated colour sensor for real time luminance calibration). Stimuli were displayed using Windows (version 10) and Matlab (R2021a), in conjunction with routines from Psychtoolbox 3 (Brainard, 1997; Kleiner et al., 2007; Pelli, 1997). Testing took place in darkened rooms, where participants viewed the displays from approximately 60cm while making responses via a keyboard.

### Design: Experiment 1, Discrimination Tasks

Prior to the discrimination tasks, participants were shown physical pieces of coloured paper and coloured cellophane, as well as an array of example complex stimuli of different filter transmittances and surface reflectances. Surface reflectance was explained as equivalent to the colour of the paint on the dragon (or the colour of the paper), which remain the same even when the filter changed (or the paper is viewed through coloured cellophane). Tristimulus value or ‘local colour’ was explained as the pixel value over the dragon part of the scene, or the combined colour of the paper and cellophane. The experimenter used a piece of black cardboard with a small, central aperture to show the local tristimulus coordinates of a small part of each example dragon when viewed without context, to demonstrate that pixels could be grey when surface and filter colours were opposite. Participants were also shown examples of the ‘reduced’ stimuli and were instructed to think of these as simulated scenes where a coloured cardboard cut-out of a dragon is against a background of another piece of coloured cardboard, where the entire scene is viewed through a filter like those in the complex scene. When making judgements of surface reflectance in the reduced stimuli, participants were instructed to judge the colour of the cardboard from which the dragon cut-out was made. We used this description of the reduced stimulus as simulating a flat cardboard dragon to help participants understand how they could interpret the reduced stimulus as a physical scene in a manner analogous to the complex stimulus when performing the surface reflectance task. After these instructions, participants completed a practice block of 12 trials while the experimenter was present before proceeding.

Across different blocks of the colour discrimination task, participants were shown either complex, reduced, or single-colour stimuli and were instructed whether they were selecting the odd-one-out in terms of surface reflectance properties (surface task) or ‘local colour’ (tristimulus value). On each trial, participants viewed 4 stimuli sequentially, using a keypress to cycle through the 4 options; there were always 4 different filter colours across the options. Stimuli were 14 degrees visual angle wide and 8 degrees high, and were presented centrally against a black background, with 200ms black screen between stimuli. Including a brief black screen at the time of the transition between the stimuli makes it less likely that the simulated surface changes would ‘pop-out’ as in an abrupt transition from one stimulus to the next, which can improve colour constancy (Foster et al., 2001). Participants were instructed to cycle through the options as many times as needed, then select the singleton using the numbers 1 to 4 (each stimulus was numbered below the image). The singleton could only be selected after all 4 stimuli had been viewed at least once. After response selection, participants were given immediate feedback on their accuracy (‘CORRECT’ or ‘WRONG’ was displayed in white text on a black screen for 1s) then the next trial began.

In the surface reflectance blocks, each trial included 3 stimuli with the same surface reflectance (one of the four base surface reflectances in Figure 1**B**, left panel), and a singleton that was either 18, 30, 60, 90 or 180 degrees different in hue. Within each trial, the 4 stimuli had 4 different filters (the 4 filters in Figure 1**B**, middle panel). Similarly in the tristimulus task, 3 stimuli had the same tristimulus coordinates (one of the four base tristimulus coordinates in Figure 1**C**, right panel). As for the surface task, in the tristimulus task, on each trial the 4 stimuli had different filters, but there was a different set of 4 filters for each base tristimulus value, as shown in Figure 1**C**. For the 3 distractor stimuli, 3 different surface reflectances were selected so that their resultant tristimulus coordinates would match the base value. The remaining filter (singleton stimulus) was paired with a surface reflectance was either 18, 30, 60, 90 or 180 degrees different in surface reflectance hue from the surface that would have matched the other stimuli in tristimulus coordinates. In this way, we sought to use an equivalent range of singleton differences for the surface and tristimulus tasks.

In each block, there were trials of each combination of the 4 relevant base colours (surface or tristimulus coordinates), where the singleton was under each of 4 relevant filters, and differed from the distractors at one of the 9 levels of surface reflectance hue difference (18, 30, 60 or 90 degrees either clockwise or counter-clockwise, or 180 degrees), giving 4x4x9=144 trials per block of trials. In each of two sessions, participants completed one block of each stimulus (complex or reduced) and task (surface or tristimulus), and a block of the single-colour stimulus (tristimulus task only); 10 blocks in total. The order of these blocks was counter-balanced across participants.

For the surface reflectance task, we considered each participant’s accuracy as a function of the singleton’s difference from the distractors in surface reflectance hue angle. For the tristimulus match task, we instead considered accuracy as a function of the measured **a*b\*** values: for each trial, we used the singleton’s measured Cartesian distance from the other 3 stimuli in the **a*b\*** plane. For both tasks, the empirically measured tristimulus coordinates of the target dragon varied systematically with surface reflectances of different hue angles. The stimuli that were offset by 18, 30, 60, 90 or 180 degrees had Cartesian distances in the **a*b\*** plane (from their base stimulus) with averages of 11.3, 17.7, 34.0, 47.6 and 67.3 respectively (surface task stimuli) and 11.3, 17.2, 33.6, 48.3 and 69.7 respectively (tristimulus task stimuli).

### Data Analysis: Experiment 1, Discrimination Tasks

Individual participants’ accuracy across hue difference in each colour discrimination task was fit using a log Weibull psychometric function (Kingdom, 2016, p. 80), given by Equation 1:

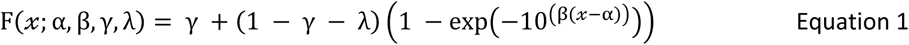

Where F(𝓍; α, β, γ, λ) is the proportion correct as a function of singleton difference from the distractors (𝓍). For the surface reflectance task, singleton difference was defined as hue angle (5 unique values). For the tristimulus coordinates task, singleton difference was defined as observed Cartesian distance in the a*b* plane, binned into 5 logarithmically spaced values. The lapse rate was set to 5% (λ=0.05), and chance rate was set to 25% (γ=0.25). We used the Matlab function *fminunc* to find the best-fitting values of the threshold (α) and slope (β) and defined the participant’s sensitivity as the inverse of the singleton difference where this function reached accuracy of 60%. To enable comparison in sensitivity across the two task types, we scaled the sensitivity values so that they were of equivalent, arbitrary units, using the average mapping between surface reflectance hue angle difference and the measured tristimulus coordinate difference to equate these measures. Statistical analyses of the sensitivity values were performed in JASP (v0.16.3, https://jasp-stats.org/), including a Bayesian repeated-measures ANOVA (van den Bergh et al., 2023).

### Design: Experiment 2, Visual Search

Each participant took part in 3 visual search tasks, completing 4 blocks of 160 trials for each task: a total of 1,920 trials across the session. On each trial participants reported, as quickly as possible, whether a target dragon colour was present or absent amongst an array of up to 16 scenes. In the first and third tasks, illustrated in Figure 3**A**, participants searched for a particular dragon surface reflectance and the scenes were made up of dragon stimuli with the colour characteristics plotted in Figure 1**B**; that is, 16 unique combinations of surface and filter properties, with only 4 unique surface reflectances. On any given block the target dragon surface reflectance was the same for all trials, and across the 4 blocks in tasks 1 and 3, participants searched for 4 different target dragon surface reflectances: red, yellow, green or blue, with block order counterbalanced across participants. On the second task, illustrated in Figure 3**B**, participants searched for a particular dragon ‘local colour’ (tristimulus value) with one target tristimulus value per block. In this task, each scene was made up of dragon stimuli with the colour characteristics shown in Figure 1**C**; that is, 16 unique combinations of surface and filter properties, with only 4 unique tristimulus values. Across blocks, participants searched for dragons of orange, purple, teal or lime in their tristimulus values, and again the block order was counterbalanced across participants. The third task was equivalent to the first, except that in task 3 only, all scenes within a trial had the same filter colour (although filter colour varied between trials). Distractor items were scenes with dragons of the 3 remaining surface reflectances (tasks 1 and 3) or tristimulus values (task 2). Due to the range of filter colours used (Figure 1**B-C**, middle column), our stimuli included scenes with different dragon surface reflectances but the same tristimulus coordinates, as well as stimuli with the same dragon surface reflectance but different tristimulus coordinates. Surface reflectance search trials contained at most 9 unique dragon tristimulus coordinates and tristimulus search trials contained up to 11 surface reflectances, as shown in Figure 1.

**Figure 3:**
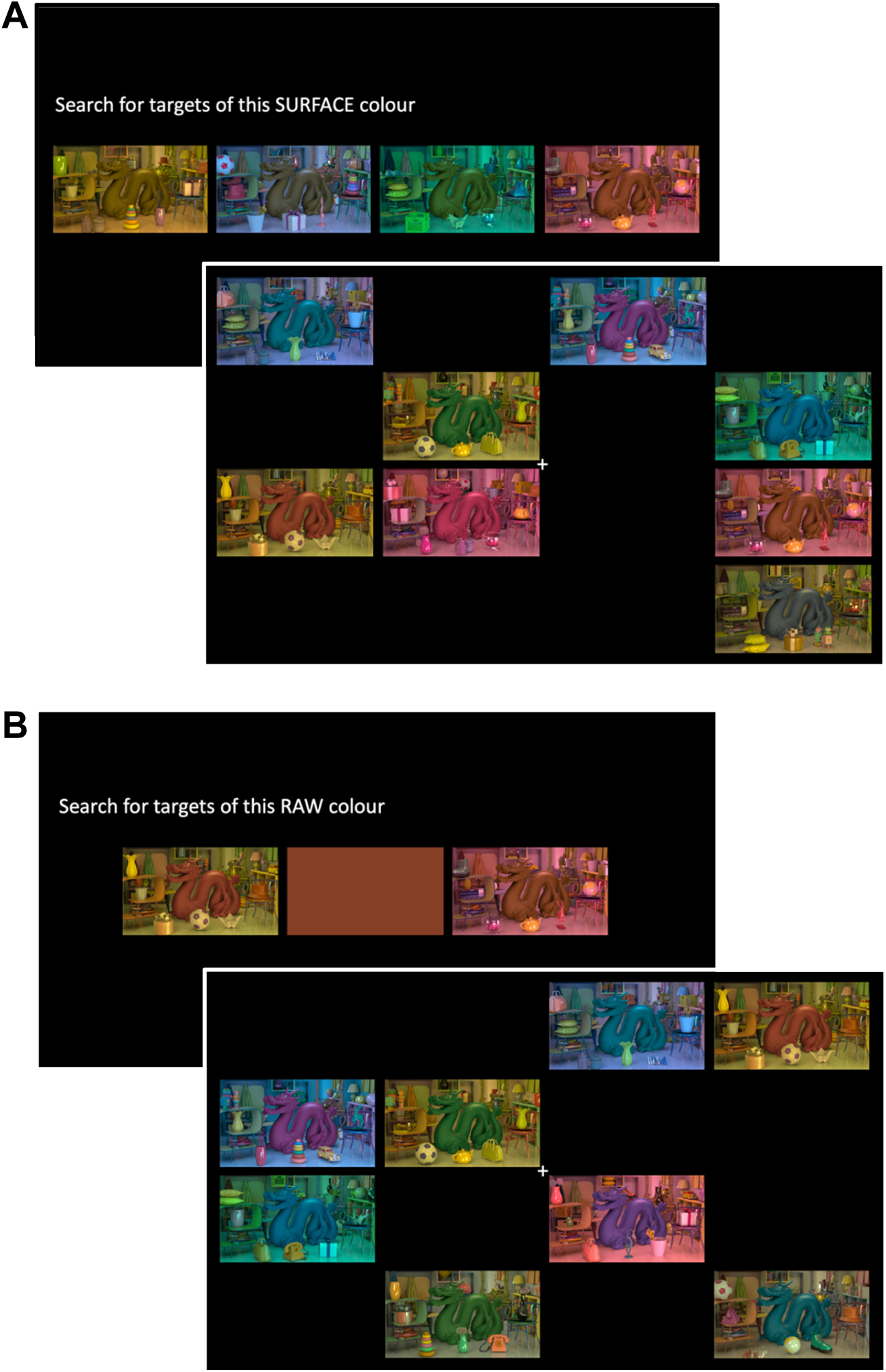
Visual Search Tasks. These schematics show an example instruction screen (upper) and target present trial (lower) for both the surface reflectance search (**A**) and the ‘local colour’ (tristimulus value) search (**B**). In the surface reflectance search, the instruction screen included examples of the target surface reflectance (e.g. yellow in the example above) under each of the four filter conditions. In the ‘local colour’ (tristimulus value) search, the instruction screen included a patch of the tristimulus coordinates of the target (e.g. orange in the example above), along with two example scenes. Both **A** and **B** show example ‘target present’ trials made up of 8 elements; across trials, there were either 2, 4, 8 or 16 elements in each stimulus.

Blocks from different tasks were interleaved, and their order was counterbalanced across participants. Each block commenced with instructions on which target surface or tristimulus value participants should search for in that block. As shown in Figure 3, for surface reflectance searches, the instruction was accompanied by examples of all 4 target stimuli, while for tristimulus search blocks the instruction was accompanied by 2 example target stimuli and a uniform patch with the average tristimulus coordinates of the targets. Within a block, after every 10 trials, participants were reminded of the target with the same instruction screen, which also provided an opportunity to take a break before they proceeded with the next trial via a button-press. On each trial, a central fixation marker (small white circle, 0.1 deg diameter) appeared 500ms before the stimulus array, and remained throughout the trial. We used the onset of the fixation marker to indicate that the array was about to appear, and we asked participants to fixate on the marker at the start of each trial, but we did not enforce fixation or track eye movements. All scenes in the stimulus appeared simultaneously and remained on the screen until the participant responded with a keypress whether a target dragon was ‘present’ or ‘absent’. After their response, participants received 1.5s of feedback on their accuracy (‘CORRECT’ or ‘WRONG’), and an update on their ‘points’ for the block: we used a points tally to encourage participants to complete each search as quickly as possible while maintaining almost perfect accuracy (minimum 97.5%). Points were added for a correct response (maximum 100, which reduced with increasing reaction time) and penalised for an incorrect response (-1000). Participants were aware that the points did not translate into any outcome and were included only for motivation. After feedback, the next trial commenced. If a participant made more than 4 errors within a block of trials, then the block aborted early and was repeated at the end of the sequence: across 12 blocks participants required an average of 2.2 (std dev 4.1) extra blocks. By requiring almost perfect accuracy, we sought to avoid participants adopting a strategy of prioritizing speed at the expense of accuracy.

The stimulus array was made up of 16 locations arranged in a 4x4 grid, centred on fixation. The total area was 40 deg wide and 22.5 deg high; individual scenes were 9.1 deg wide and 5.1 deg high, with 0.9 deg gaps between scenes. Of the 16 possible locations, between 2 and 16 contained a stimulus on each trial, while the remaining locations, gaps between locations, and the remainder of the screen were black. Each block contained 20 target-present (each containing a single target) and 20 target-absent trials, for each number of elements (2, 4, 8, or 16), presented in a randomly interleaved order. For each trial, the individual scenes were placed at random locations, without replacement, meaning that when there were fewer than 16 scenes the empty locations varied across trials.

### Data Analysis: Experiment 2, Visual Search

We analysed median reaction times for correct trials across included blocks (excluding data from any blocks that were aborted early due to more than 4 errors). For each participant, we found the search slopes using the Matlab function *polyval* to find the line of best fit between logarithmically spaced element numbers (*x*) and median reaction time (RT), i.e. RT(2*^x^*) = *a* + *mx*, where *a* is an offset. We performed statistical analyses of the slope values in JASP (v0.16.3, https://jasp-stats.org/), including a Bayesian repeated-measures ANOVA (van den Bergh et al., 2023).

### Model predictions: simplified guided search

Since our stimuli did not perfectly separate dragon surface reflectance and tristimulus values, we used a modelling process to formalise our predictions for reactions times if the dragon’s surface reflectance but not its tristimulus value were available preattentively, or vice versa. We also considered the scenario where both the dragon’s surface reflectance and tristimulus value were not available preattentively, but the tristimulus values of the filter could preattentively guide the participants’ search. We used a simple model based on Guided Search 6.0 (GS6) (Wolfe, 2021). The main aspect of GS6 that we included in the model predictions was to assume that the search process is simultaneously serial and parallel, with preattentive feature guidance. That is, preattentive values form a priority map that guides the serial search for feature values that are not available preattentively. In each of the candidate models outlined below, we assumed that only a single feature was available preattentively, rather than considering a more complex model where multiple features had varying levels of efficiency, to avoid an under-constrained model, although more complex models could be considered in future work. Similarly, we substantially simplified other aspects of the model, since our experimental design and numbers of trials were not optimised to constrain, for example, asynchronous diffusion rates, effects of search history, or inhibition of return during search. Since we required participants to perform with near perfect accuracy, we set a constant, very conservative, quitting threshold by assuming that participants always serially searched all potential targets before selecting a ‘target absent’ response. We assumed that search proceeded without replacement, since our largest set size was relatively small (16), and on average fewer than half the elements required serial search. Instead of modelling the asynchronous diffuser, we assumed that each item requiring serial search took 250 ms to process.

We considered three candidate models, with predictions illustrated in Figure 4. In the first model, we assumed that the surface reflectances of the dragons, but not their tristimulus coordinates, are available preattentively, and that surface reflectance guided search for tristimulus value. In the second model, we assumed this pattern was reversed: that the tristimulus coordinates of the dragons are available preattentively and would guide search in the surface reflectance task. In the third model, we assumed that only the average tristimulus coordinates of the filter transmittance are available preattentively, which may occur if the dragon’s visibility were crowded by its surrounding background when participants viewed the visual search array. In each model, we predicted the order of search according to the model’s preattentive feature. For each search type, we assumed that participants would search through the items in order according to their preattentive feature’s proximity to a search template location (examples indicated by red circles in **Figure 4**), starting with the closest item. As discussed above, the tristimulus coordinates of the dragon stimuli were not directly controlled, and there were some small differences between stimuli that were categorised as tristimulus matches. We reasoned that participants were unlikely to be sensitive to these small discrepancies during visual search, and when modelling the order of search according to tristimulus coordinates we assumed that these approximate matches were perfectly aligned in tristimulus coordinates, and evenly spaced (i.e. assuming that in the surface reflectance search, the stimulus tristimulus coordinates formed an evenly spaced 3x3 grid of 9 values).

**Figure 4:**
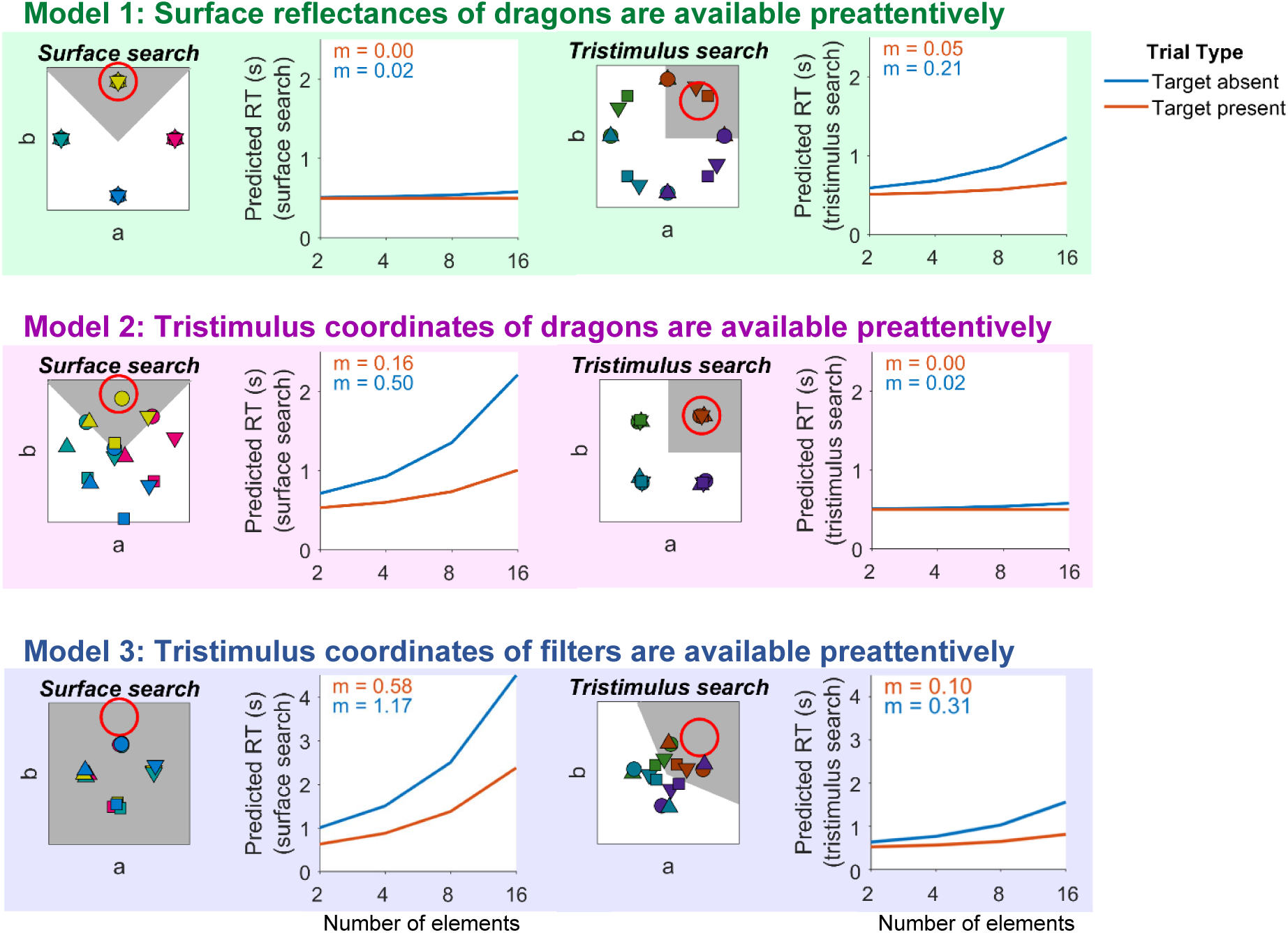
Three possible models of preattentive colour perception, including predicted visual search reaction times (RTs). Model 1 (green box, top) assumes that the surface reflectances of the dragon stimuli are available preattentively, but not their tristimulus coordinates. Model 2 (magneta box, middle) assumes that each dragon’s tristimulus coordinates, but not their surface reflectance are available preattentively. Model 3 (blue box, bottom) assumes that no features of the dragons are available preattentively, but that the tristimulus coordinates of the filters, approximating the average of each search element, can preattentively guide search. Predictions for surface reflectance and tristimulus searches are shown in left and right plots of each box respectively. To the left of each plot of predicted RTs, the preattentive colours (in CIE L*a*b* coordinates) of the 16 unique dragon stimuli are shown for the relevant model and task. Markers are coloured according to their search feature (surface reflectance for left column of plots, tristimulus coordinates for right column of plots). The red circle highlights an example search target (yellow surface or orange tristimulus coordinates), and the grey shaded region shows the region of feature values that the model assumes are searched for the example target. Predicted search slopes (m) for target present (orange) and target absent (blue) trials are indicated in the top left of each plot.

We assumed that participants could adopt a strategy optimised for our stimulus set, and only needed to check items with preattentive feature values that were potentially consistent with the target: this region is shown in the grey shaded areas in **Figure 4**. For example, assuming that surface reflectance is available preattentively, then when the participant was searching for an orange tristimulus value, only surface reflectances from yellow to red were potentially consistent with the target, whereas greenish or bluish surfaces could be rejected without scrutiny. Note that the tristimulus coordinates of the filters are uninformative about the dragon’s surface reflectance, so the only case where there is no preattentive guidance of attention is in the surface reflectance search in Model 3: in this case the model predicts that both target absent and target present searches will be inefficient (with a search slope in the target absent trials approximately twice that of the target present trials).

For each model, search was assumed to terminate in the target-present trials once the target item was searched. If the target was absent, search continued until all items that were potentially targets (items in the grey shaded area) were scrutinised and rejected. We assumed a very brief but non-zero time of 5ms per item that was rejected preattentively, based on observations that reaction times in basic feature search are not perfectly parallel, but increase slightly with the log of the set size (Lleras et al., 2020; Wang et al., 2018).

Predictions of search times (in s) for target absent (ST_absent_) and target present (ST_present_) searches, as a function of the number of items (*n*), are defined by Equations 2-3:

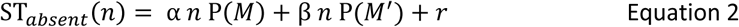

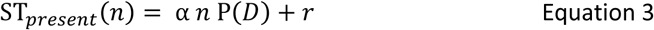

Where α is the time taken to process an item requiring serial search (fixed at α = 0.25), β is the time taken to reject a distractor requiring parallel search (fixed at β = 0.05) and *r* is the time taken to generate a motor response based on the participant decision (fixed at 𝑟 = 0.5).

Across models, the only variables that changed were P(𝑀), which is the probability that a distractor matches the search template in a target absent search, P(𝑀′), defined as P(𝑀′) = 1 − P(𝑀), and P(𝐷), the probability, in a target present search, that a distractor is closer to the template than the target. The probabilities used for each model and search type are shown in **Table 1**.

**Table 1:** Probabilities used in modelling. For both search types, there were 4 possible targets (with a maximum of 1 on each trial), and 12 possible distractors. In each case, P(𝑀) is the proportion of the 12 distractors that are in the search region (grey shaded areas in Figure 4). In some cases, P(𝐷) varies depending on which of the 4 targets is present, and on whether the distractor is closer to the search template than the target (and so the probability it would be selected over the target is 1) or is the same distance from the search template (where there is a 1 in 2 chance it will be selected before the target). To show how these probabilities were derived, for P(𝐷) we express each probability as the number of targets (out of 4) and number of distractors (out of 12) matching each scenario.

| Model | Surface reflectance search task |  | Tristimulus search task |  |
| --- | --- | --- | --- | --- |
| | $P(M)$ | $P(D)$ | $P(M)$ | $P(D)$ |
| 1 | $\frac{0}{12}$ | $\frac{4}{4} \times \frac{0}{12} = 0$ | $\frac{0}{12}$ | $\frac{2}{4} \times \frac{2}{12} \times \frac{1}{2} = 0.0417$ |
| 2 | $\frac{5}{12}$ | $\frac{2}{4} \times \frac{2}{12} \times \frac{1}{2} + \frac{1}{4} \left( \frac{2}{12} + \frac{3}{12} \times \frac{1}{2} \right) = 0.114$ | $\frac{0}{12}$ | $\frac{4}{4} \times \frac{0}{12} = 0$ |
| 3 | $\frac{12}{12}$ | $\frac{4}{4} \times \frac{12}{12} \times \frac{1}{2} = 0.5$ | $\frac{3}{12}$ | $\frac{4}{4} \times \frac{2}{12} \times \frac{1}{2} = 0.0833$ |

Intuitively, P(𝐷) can be understood by considering an example: in Model 2, where the dragon’s tristimulus coordinates are available preattentively, then we assume that the dragon’s tristimulus coordinates guide the order in which items are searched when searching for a target surface reflectance. If participants are searching for a yellow surface reflectance (as in the example in **Figure 4**), then, statistically, if a target is present, it is likely to be the item in the array with a tristimulus coordinates that are closest to yellow, and so will be the first item searched. Of the 4 possible yellow surface target stimuli (shown as yellow symbols in **Figure 4**, Model 2, left), 1 target (circle) will always have the most yellow tristimulus coordinates of any in the array when present, 2 targets (triangles) will have the most yellow tristimulus coordinates in the array, but may be amongst at most 2 distractor stimuli of the same yellow tristimulus coordinates, and only the final target (square) could potentially be present along with at most 2 distractor stimuli that are closer to yellow in their tristimulus coordinates.

Although predicted RTs vary with model parameters, in all cases Model 1 predicts higher RT slopes for tristimulus coordinates search than for surface reflectance search, whereas Models 2-3 predict the reverse pattern. Model 3 predicts longer RTs than Models 1-2 with the same parameters (note the different y-axis scaling for Model 3 in **Figure 4**), although shorter RTs can be modelled by adjusting the free parameters α, β, and 𝑟. Across model parameters, the most reliable difference between Models 2 and 3 is in the relative search slopes for target absent versus target present trials in the surface reflectance search: since Model 3 assumes there is no preattentive guidance of this search, the search slope for target absent trials is predicted to be twice that of the target present trials. In Model 2, which predicts preattentive guidance of attention by the dragon’s tristimulus coordinates, the target absent trials are predicted to have a search slope more than twice the target present trials: for the parameters in **Figure 4**, the target absent search slope is 3.7 times greater than the target present slope.

## Results

### Experiment 1: Colour constancy and tristimulus coordinates discrimination thresholds

To measure participants’ colour constancy for the dragon objects when the dragon is attended, we used a 4-interval-force-choice (4IFC) discrimination task to identify the smallest surface reflectance difference that was required for participants to reliably identify the odd-one-out in dragon surface reflectance. In the 4IFC design, participants viewed the stimuli sequentially with brief black screens between stimuli, to ensure that discrimination thresholds reflected performance when the dragons were individually attended, but without the potential for surface changes to ‘pop-out’ as in abrupt stimulus transitions without a transient (Foster et al., 2001). We used a separate 4IFC task, with an analogous design, to identify how reliably participants could identify the odd-one-out in dragon tristimulus value.

Figure 5 shows discrimination thresholds for detecting changes in surface reflectance and changes in tristimulus values across two types of stimuli: complex stimuli (those used in the visual search task, below) and reduced stimuli (simplified versions of the complex stimuli that removed many cues to colour constancy). For tristimulus coordinates we also included a baseline measure of discrimination thresholds for uniform patches (the ‘single colour’ condition). For the complex stimuli used in the visual search task, sensitivity to small differences in tristimulus value or surface reflectance was very similar. Sensitivity to surface reflectance differences was slightly higher than for tristimulus values (i.e. discriminating surface reflectance was slightly easier), but the difference was not significant (see Figure 5B-C, and below for statistical tests). When the singleton varied by a larger difference in surface or tristimulus coordinates, performance reached similarly high levels of accuracy of 93.5% (surface reflectance) and 93.0% (tristimulus coordinates) for the complex stimuli of 90 degrees hue difference (see purple lines in Figure 5A).

**Figure 5:**
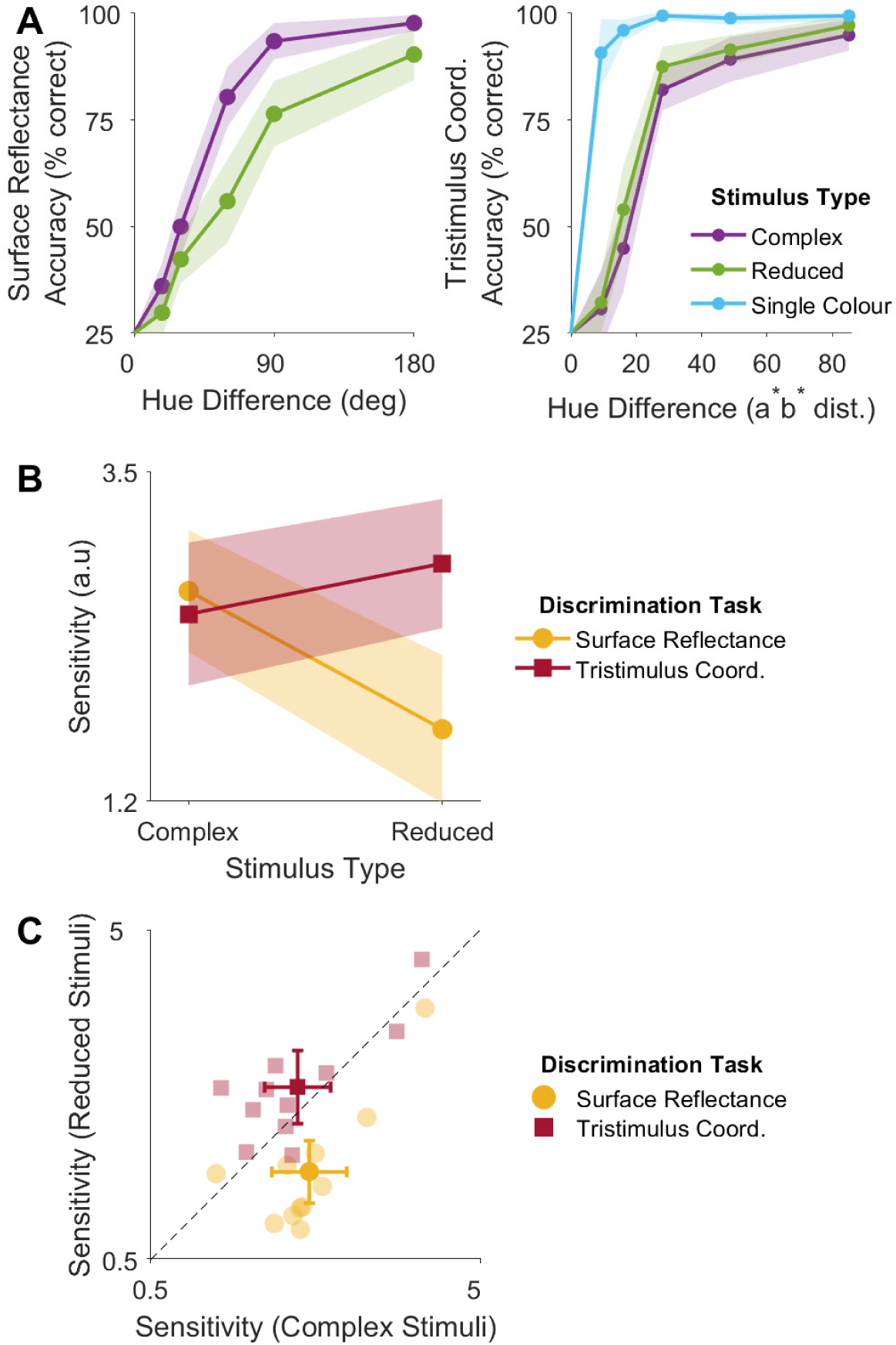
Discrimination thresholds. Average accuracy in the 4IFC tasks based on surface reflectance (**A**, left) and tristimulus coordinates (**A**, right). Each individual’s data were fit with a psychometric (log Weibull) function for each condition, with sensitivity defined as 1/hue difference threshold for 60% accuracy. Sensitivity across condition and the complex and reduced stimulus types is shown in **B** (averages) and **C** (individual sensitivity values). Error bars are 95% confidence intervals of the between-subject means (n=11).

When many image cues were removed in the reduced stimuli, sensitivity to surface reflectance decreased and sensitivity to tristimulus coordinates increased. A 2x2 Bayesian repeated measures ANOVA of the effect of discrimination type (surface reflectance or tristimulus coordinates) and stimulus complexity (complex or reduced) on sensitivity revealing strong evidence in favour of a main effect of discrimination type (BF_incl_=425.5), strong evidence in favour of a main effect of stimulus complexity (BF_incl_=358.7), and strong evidence in favour of an interaction between discrimination type and stimulus complexity (BF_incl_=1,264.5). Simple main effects showed there was anecdotal evidence against a difference in sensitivity between discrimination types for the complex stimuli used in the visual search task (BF_10_=0.327), (note that this shows that threshold discriminability was closely matched for the conditions used in the two visual search tasks, below), but strong evidence that sensitivity was higher when discriminating changes in tristimulus coordinates rather than surface reflectance for the reduced stimuli (BF_10_=13.02).

Simple main effects of stimulus complexity on sensitivity revealed strong evidence of a difference for the surface reflectance task (BF_10_=157.5) and anecdotal evidence of a difference for the tristimulus value task (BF_10_=1.564). This is seen in Figure 5C, where red squares tend to lie above the diagonal and yellow circles tend to lie below. That is, sensitivity to tristimulus coordinates tended to improve as sensitivity to surface reflectance decreased. For the tristimulus coordinates task we also included a ‘single colour’ condition to estimate the extent to which discrimination sensitivity in the tristimulus coordinates task might have been reduced by small stimulus variations arising from the way in which the stimuli were generated (see Methods). Accuracy for discriminating tristimulus coordinates in the ‘single colour’ condition is shown in Figure 5A (right plot, blue line). In this condition, there was strong evidence that discrimination sensitivity was higher than in the ‘reduced’ condition (Bayesian paired *t*-test: BF_10_=31.833), suggesting that the stimulus variations in tristimulus coordinates are unlikely to have limited sensitivity in the tristimulus tasks for the complex and reduced stimuli.

### Experiment 2: Visual Search

Next, we compared how efficiently tristimulus value and surface reflectance could guide search for a target dragon. In all the visual search tasks we used only the complex stimuli, for which participants were more colour constant, and had approximately the same sensitivity for the dragon’s tristimulus value and its surface reflectance. Participants searched for a target surface reflectance or a target tristimulus values, completing different search types and searching for different target colours across different blocks of trials. On each visual search trial, participants saw an array of up to 16 elements, where each element was a scene stimulus containing a central dragon object (see Figure 3).

Figure 6 shows the median reaction times across each task. We found that search was clearly less efficient when searching for surface reflectance than for tristimulus value. As the number of elements increased, reaction time increased more steeply for the surface reflectance search (Figure 6A) than for the tristimulus value search (Figure 6B). The direction of this effect was very consistent; across every participant (n=10) the search slope for the surface reflectance search was higher than for the tristimulus value search in both target present and target absent trials (Figure 6D). A 2x2 Bayesian repeated measures ANOVA of the effect of search type (surface or tristimulus value) and trial type (target absent or present) on search slope revealed strong evidence of main effects of search type (BF_incl_=36.866), and trial type (BF_incl_=92.277), and extremely strong evidence of a significant interaction between these factors (BF_incl_=9995.038).

**Figure 6:**
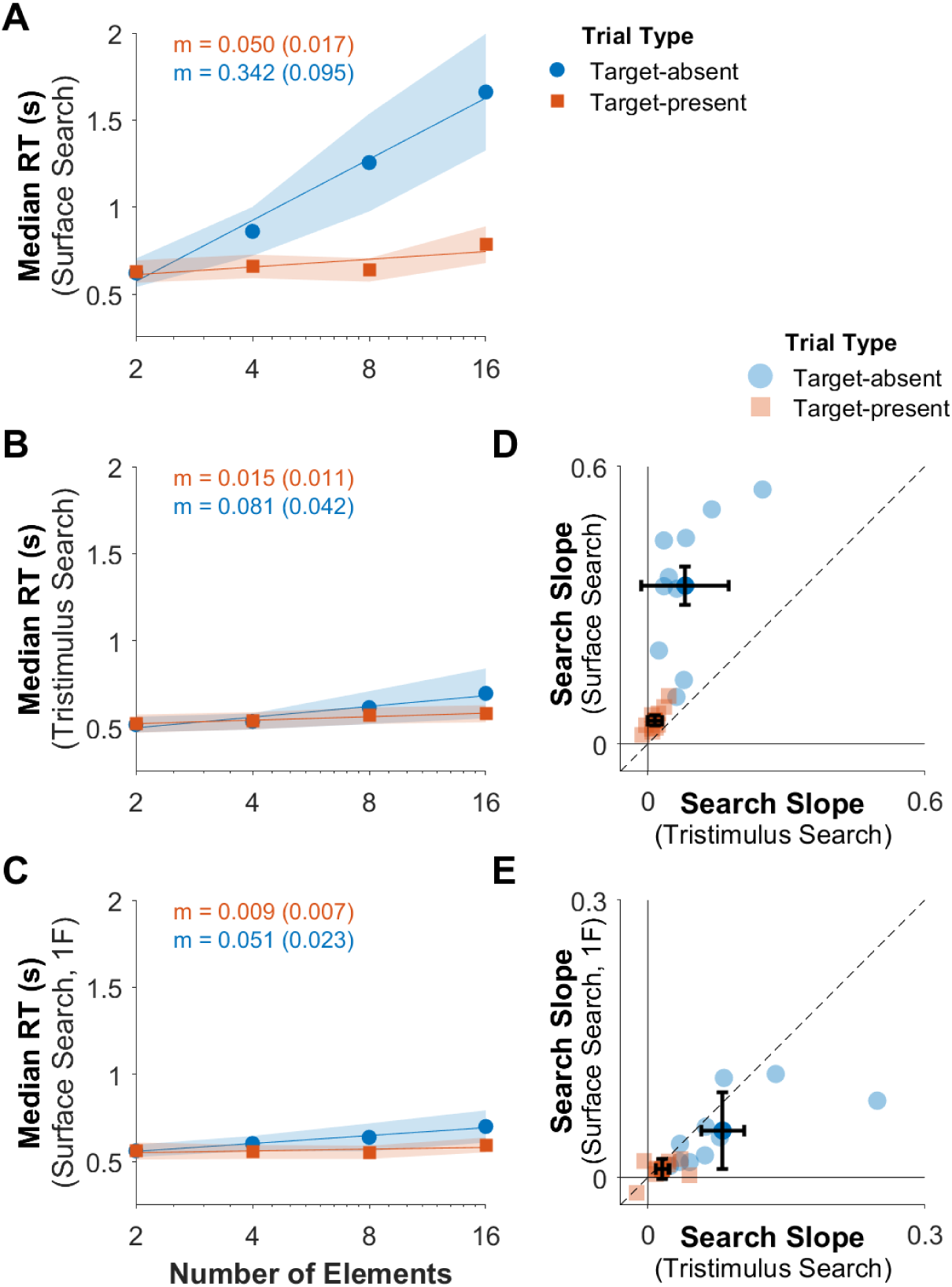
Median reaction times (n=10) for the surface reflectance search (**A**), tristimulus value search (**B**), and surface reflectance search where each trial contained elements under the same filter (**C**). Shaded error bars are 95% confidence intervals of median RTs, and lines show the line of best fit to the median data. Text insets indicate the mean slope (m) for fits to individual data (and 95% confidence intervals); see Methods for details. **D** and **E** show search slopes for individual participants across tasks, and across target absent and target present trials, along with the group averages (at centres of error bars indicating 95% confidence intervals), corresponding to the data in the text insets of **A-C**. All participants met strict accuracy criteria (minimum 97.5% correct for each search type). Mean accuracies (with 95% confidence intervals) were slightly higher for ‘target absent trials’: 99.4% (0.25%), 99.4% (0.30%) and 99.3% (0.23%) for the tasks in **A**, **B**, and **C**, respectively, than for ‘target present trials’: 97.4% (0.46%), 98.1% (0.40%) and 98.4% (0.41%) for **A**, **B** and **C**, respectively.

Typically, in serial or ‘unguided’ search, search slopes for target present trials are expected to be half that of target absent trials, since on target present trials participants will, on average, serially search half the items before finding the target (e.g. Wolfe, 2021). However, in the surface reflectance search (Figure 6A), slopes for target absent trials were at least 5 times that of the target present trials for every participant, with an average 6.10-fold increase relative to the target-present trials. This suggests there was some preattentive guidance of visual search in the surface task, as well as in the other two search types.

Although our stimuli partly dissociate the surface reflectance and tristimulus coordinates of the dragons, these attributes were not fully independent in the visual search stimuli. Because of this, if either the dragons’ surface reflectance or their tristimulus coordinates are available preattentively, then this could guide search for the other feature. If the tristimulus coordinates of the filter but not the dragon were available preattentively (as might occur if the dragon’s visibility were crowded by its surround when the scenes were viewed in the visual search array) then preattentive perception of the filters’ tristimulus coordinates could guide the tristimulus but not the surface reflectance search. To estimate the expected effect of this guidance on search slopes, we implemented a reduced version of the Guided Search 6.0 (GS6) model (Wolfe, 2021; see Methods for details), assuming either that the dragons’ surface reflectances are available preattentively while their tristimulus coordinates are not (Model 1, green box in Figure 4), that the dragons’ tristimulus coordinates are available preattentively while their surface reflectances are not (Model 2, magenta box in Figure 4), or that only the tristimulus coordinates of the filters are available preattentively (Model 3, blue box in Figure 4).

Adjusting the time constants of the model changes the predicted reaction times, but in every case Model 1 predicts that surface reflectance search is more efficient than tristimulus search and Models 2-3 predicts the reverse: in this way, our results are inconsistent with Model 1. Model 3 assumed that the only stimulus feature available preattentively was the tristimulus coordinates of the filter, which in our design was uninformative regarding the dragon’s surface reflectance, so this Model cannot account for the evidence of preattentive guidance seen in our data for the surface reflectance search (Figure 6A), where search slopes for target absent trials were more than twice those for the target present trials. From this, our data are most consistent with the dragons’ tristimulus values, but not their perceived surface reflectances, being able to preattentively guide visual search, and the descriptive model in Model 2 provides a reasonable account of the observed reaction times.

In the final visual search task (Figure 6C), participants were searching for a target surface reflectance (as in the first task, shown in Figure 6A) but on each trial, all elements were scenes with a simulated filter of the same colour. In this case, search was highly efficient, and search slopes tended to be lower than for the tristimulus coordinates search (Figure 6E), although there was minimal evidence of an effect: a 2x2 repeated measures ANOVA of the effect of search type (surface reflectance under single filter or tristimulus coordinates) and trial type (target absent or present) on search slope no evidence of a main effect of search type (BF_incl_=1.043), strong evidence of an effect of trial type (BF_incl_=12.945), and anecdotal evidence against an interaction (BF_incl_=0.863). There are at least two possible accounts of high search efficiency in this third task; first, efficient search could result from preattentive dragon tristimulus coordinates (Model 2), with increased efficiency from greater distractor homogeneity relative to the surface reflectance search with heterogenous filters (Wolfe, 2021); alternatively, when the filters were homogenous across all elements in the array, the perceptual scission of all elements may occur simultaneously across elements, allowing participants to search for the target surface reflectance efficiently.

## Discussion

We used visual search as a tool to test whether the perception of surface reflectance and tristimulus coordinates require attention, or are available preattentively, to provide new insight into the neural mechanisms underlying colour constancy. Our results suggest that tristimulus values, but not surface reflectances, are available preattentively when there are multiple filter conditions across the visual field.

### Surface and tristimulus coordinates thresholds were similar for complex stimuli

To maximally dissociate the target surface reflectance and tristimulus values in our stimuli, the target (central dragon object) always had a saturated surface reflectance, and each scene was viewed through a simulated translucent filter, also of a saturated colour (see Methods). Since these stimuli included saturated translucent filters, we expected that colour constancy would be imperfect for these stimuli, since colour constancy is known to decrease with the saturation of a simulated illuminant (Reeves & Amano, 2020). Conversely, since the tristimulus values are of less behavioural relevance and may not be represented after perceptual scission of surface and illuminant (or filter) properties, we also expected participants may be imperfect in judging tristimulus coordinates. The discrimination thresholds (Experiment 1, Figure 5) confirmed that participants were below ceiling performance in both these tasks.

Importantly, for the visual search experiment, we did not require that participants had perfect accuracy in identifying either the dragon’s surface reflectance or its tristimulus value. Instead, we sought to satisfy two requirements: first, that participants were comparably accurate when identifying surface and tristimulus coordinates of the dragons when the stimuli were attended individually. The similar discrimination sensitivity values for tristimulus values and surface reflectance (for complex stimuli) show that this requirement was met. Second, to ensure the visual search stimuli in both tasks were of sufficiently large difference along the relevant feature dimension (surface reflectance or tristimulus value) that participants were approaching ceiling accuracy. This was to ensure that if surface and/or tristimulus coordinates were able to preattentively guide search, then there would be sufficient feature difference for efficient search (Carter, 1982; Duncan & Humphreys, 1989; Nagy & Sanchez, 1990). Discrimination thresholds show that this requirement was also met, with performance approaching ceiling level for the colour differences used in the visual search tasks (90 or 180 degrees hue difference in surface reflectance, or an equivalent change in tristimulus value).

The discrimination thresholds for surface reflectance and tristimulus value changes also provide insight into whether any colour constancy for the complex stimuli is based on a change in the perceived hue of the dragon, or could be attributed entirely to explicit reasoning (as discussed in Radonjic & Brainard, 2016), without any shift in perceptual experience. To test this, we compared how judgements of surface reflectance and tristimulus value changed when image cues relevant to colour constancy were removed. As expected, sensitivity to surface reflectance was lower in the ‘reduced’ condition than for the complex stimuli, even though the complex and reduced stimuli were closely matched in their tristimulus coordinates (Radonjic et al., 2015). If the improved colour constancy for complex stimuli is at least partly based on a shift in the perceived colour, and the percept of surface reflectance and filter properties replaces the representation of tristimulus coordinates, then when colour constancy decreases, sensitivity to tristimulus coordinates should improve. Using Bayesian statistics, we compared the strength of evidence in favour of no effect of stimulus condition with the evidence in favour of a consistent difference. Our data did not clearly differentiate between these alternatives, but provided anecdotal support in favour of a sensitivity to tristimulus coordinates being higher for the reduced stimulus, consistent with perceptual experience shifting towards greater colour constancy as complexity increased.

### Tristimulus values, but not surface reflectance, can preattentively guide search under heterogenous filters

Having established that participants were of similar accuracy for discriminating surface reflectance or tristimulus value changes for these stimuli, in Experiment 2 we used visual search to test whether either surface reflectance or tristimulus values of the dragons were available preattentively. Although the surface reflectance and tristimulus values of the targets were not fully dissociated, we selected the scene elements so that in both search tasks the dragons’ tristimulus coordinates did not uniquely predict their surface reflectance and vice versa. Across every participant, for both target present and target absent trials, search slopes were higher for searches based on surface reflectance than those based on local tristimulus value, and the search slopes for target absent trials in the surface reflectance search task were at least 5 times greater than for target present trials, consistent with some preattentive guidance towards the targets when they were present. Using a reduced version of the Guided Search 6.0 (GS6) model (Wolfe, 2021), we found that our data are well described by a model assuming that the tristimulus coordinates of the dragons guides search in both tasks, but that when there are heterogenous filters across the scene, perceived dragon surface reflectance is only available when the item is attended.

The control condition, where participants searched for surface reflectance under a single filter, showed that search for surface reflectance can be efficient when all elements are under a homogenous filter. In this condition participants could be preattentively searching for a tristimulus value relative to the average colour of all elements, with better efficiency than in the main surface reflectance search task caused by the increased homogeneity of the distractors (Wolfe, 2021). Note that the common filter changed across trials, so this strategy is not identical to searching for a specific set of tristimulus coordinates, since, unlike in the tristimulus search task, the participants could not know the target tristimulus value before each trial commenced. Alternatively, for displays with a common filter across all scene elements, perceptual scission may occur simultaneously across the entire display, so that the surface and filter properties are perceptually separated simultaneously for all elements, allowing for an efficient search for surface reflectance. This would imply that scission can be performed over a more global spatial scale when the filter conditions are uniform, but that attention to each element was required to perceive surface reflectance when under varying filters. These possibilities may be explored in future work.

Although our stimuli varied filter transmittance properties, not illumination, we think it likely that the preattentive perception of tristimulus coordinates, but not surface reflectance, would generalise to other instances requiring perceptual scission of surface reflectance properties from other factors, as in surface colour constancy across varying illumination. Perceiving surface properties through a translucent filter affects tristimulus values in ways that not identical to changing the spectral properties of the illumination (e.g. Faul & Ekroll, 2002). However, perceptually separating surface properties from those of a filter or illuminant is conceptually similar in that both require scission of colocalised information, resulting in perceptual ‘layers’, and the questions of how these computations are implemented by the visual system have been considered in analogous ways (Faul & Ekroll, 2012; Gerbino et al., 1990; Khang & Zaidi, 2002; Soranzo & Gilchrist, 2019). Based on the computational similarity of these problems, we think it likely that their underlying neural processes are likely to be analogous, and so, based on our results, we expect that attention is required for colour constant perception of surface properties when scenes have spatially heterogenous illumination.

### Differences in search efficiency cannot be explained by background complexity, crowding or distractor heterogeneity

We sought to equate the surface reflectance and tristimulus value search tasks as much as possible, to provide a fair comparison of whether they differed in search slopes. Separating a search target from a background can increase search time (Wolfe et al., 2002) meaning that the scenes surrounding each dragon object, which are necessary for colour constancy, likely increased search times. The dragon was in the same location in every scene to minimise spatial uncertainty. Importantly, the scene structure did not vary between the surface reflectance and tristimulus search tasks, meaning that factors related to scene structure or the spatial layout of the stimuli cannot account for differences in search slopes across tasks.

The constant scene structure used in all elements also rules out an explanation based on crowding, or lack of acuity for peripheral elements. For instance, if the dragon objects were crowded by their scene background, the colour of the dragon may be difficult to perceive without an eye movement towards the element, which could account for a pattern of serial search. This scenario most closely aligns with model 3, which assumed that only the filter tristimulus value was available preattentively, while the dragon surface reflectances and tristimulus values were not. However, this model’s predictions could not account for the evidence of preattentive guidance of search in the surface reflectance task in our results. Furthermore, the patterns of preattentive search in the tristimulus value task and the surface reflectance under single filter were obtained for stimuli with identical spatial properties, showing that crowding and acuity did not preclude preattentive search.

Search efficiency reduces with the heterogeneity of the distractor items, and with some feature relationships between the target and the distractors (e.g. Becker, 2010). To equate our tasks for these, both surface reflectance and tristimulus value tasks had similarly heterogeneous distractor items: both tasks included 4 unique values of the target feature, and heterogeneous filter colours. The tasks were approximately matched in terms of the alternate feature: in the surface reflectance task there were at most 9 different tristimulus values, and in the tristimulus value task there were at most 11 different surface reflectances. The impact of heterogeneity depends not only by the number of unique distractors, but also on their distance to the target element. The predicted patterns of difference are captured by the two models assuming preattentive guidance by a feature of the dragons (first two models in Figure 4): the model predictions are not perfectly symmetric, with Model 2 predicting steeper search slopes in the surface reflectance task than Model 1 predicts in the tristimulus value task (see Methods). However, these differences in distractor heterogeneity do not affect the main distinction between these models’ predictions: that in Model 1, search is more efficient in the surface reflectance search, and in Model 2, search is more efficient for tristimulus value search.

### Differences in search efficiency cannot be explained by perceptual accuracy for different dimensions

At the suprathreshold levels of hue difference used in the search tasks participants were at ceiling performance in both discrimination tasks. This is consistent all participants being able to meet the strict accuracy threshold across all search tasks (minimum 97.5% accuracy for each block). Using stimuli with large, easily discriminable differences was necessary since small but discriminable colour differences cannot guide search preattentively, whereas larger differences can (Nagy & Sanchez, 1990). For this reason, both the surface reflectance and tristimulus value tasks included only 4 unique values of the target feature, equally spaced, producing large differences in the target feature values. These were sufficiently different to guide preattentive search in the tristimulus value search. In the surface reflectance search, surface reflectances were separated by an equivalent hue angle (90 degrees) and were more saturated than the 4 tristimulus values in the tristimulus search task (as seen by comparing Figure 1B, left panel with Figure 1C, right panel), meaning that in the surface reflectance task the 4 unique values were slightly more separated than in the tristimulus task. This suggests that if surface reflectance were able to preattentively guide attention, the surface task stimuli should have been sufficiently different to allow near-parallel search. In these ways, the difference in visual search slopes cannot be accounted for by differences in how accurate participants were in judging the surface and tristimulus values when the stimuli were considered individually, nor by the degree of feature separation along the relevant dimension, across the surface reflectance and tristimulus value searches.

### Interpretation, and relation to previous work on the timescale of colour constancy

Overall, our results are most consistent with the visual system decomposing tristimulus values into surface and filter (or illuminant) properties via a process that requires attention and takes time to occur. In previous work, the timescale of colour constancy has been considered primarily in the context of the timescale of chromatic adaptation, which includes both fast (∼50ms to 1s) and slow (20-50s) components (Fairchild & Reniff, 1995; Foster, 2011; Rinner & Gegenfurtner, 2000). Neither of these effects provides a plausible account of our data: the timescale of slow chromatic adaptation means it can at most contribute to colour constancy over longer timeframes, without accounting for colour constancy over brief presentations, including the range of reactions times here. The fast component is argued to reflect primarily adaptation at the receptoral level (Rinner & Gegenfurtner, 2000), which cannot account for the incorporation of higher-level cues to colour constancy. Furthermore, if perceiving surface reflectance depended only on fast receptoral adaptation that normalises to local image statistics, then this should occur simultaneously across the visual field even when the filters are heterogenous across the visual field, allowing preattentive search after this initial normalisation.

The pattern of serial search for surface reflectance found here suggests that scission of surface and filter properties, like the binding of some feature conjunctions, requires deployment of attention (Treisman, 1996). This offers new constraints to understand how colour constancy is implemented in the brain, suggesting that when the visual field contains multiple filter or illumination conditions, scission may not automatically occur simultaneously across all the different local illumination conditions, but within localised, attended, subunits of the scene. The arrays used in our visual search task were highly artificial, but spatially non-uniform illumination also occurs frequently in natural scenes, where three dimensional objects can cast shadows and produce interreflections and lead to scenes which require local, spatially restricted computations of colour constancy (e.g. as discussed in Werner, 2014). Psychophysically, local colour constancy effects are demonstrated by studies such as Radonjic et al. (2015), who found that cues to three dimensional structure can improve colour constancy for artificial stimuli, where the simulated structure is consistent with a simple object (cube) with each of its three visible sides illuminated differently. Our results suggest that in scenes such as these, with spatially non-uniform illumination or filter conditions, these local colour constancy computations require attention. The role of attention in feature binding has been used to argue for a critical role of top-down or recurrent neural processing in the binding of simple visual features (e.g. Bouvier & Treisman, 2010; Reynolds & Desimone, 1999; Treisman, 1996; Wyatte et al., 2012); our results suggest there may be a similar role for top-down or recurrent processes in the neural mechanisms underlying colour constancy where the filter or illumination conditions are heterogenous across the scene. This is consistent with evidence that relatively complex visual cues contribute to colour constancy, including cues that depend upon the interpretation of the scene’s 3D structure (Anderson & Kim, 2009; Bloj et al., 1999; D’Zmura & Lennie, 1986), or object memory (Granzier & Gegenfurtner, 2012). Our results suggest that these computations do not take place preattentively, but may rely attentional selection.

## Acknowledgements

This project was funded by the Australian Research Council (ARC) (DP220100747 to EG). We thank Colin Clifford for helpful discussions on the manuscript.

---

**Figure S1:**
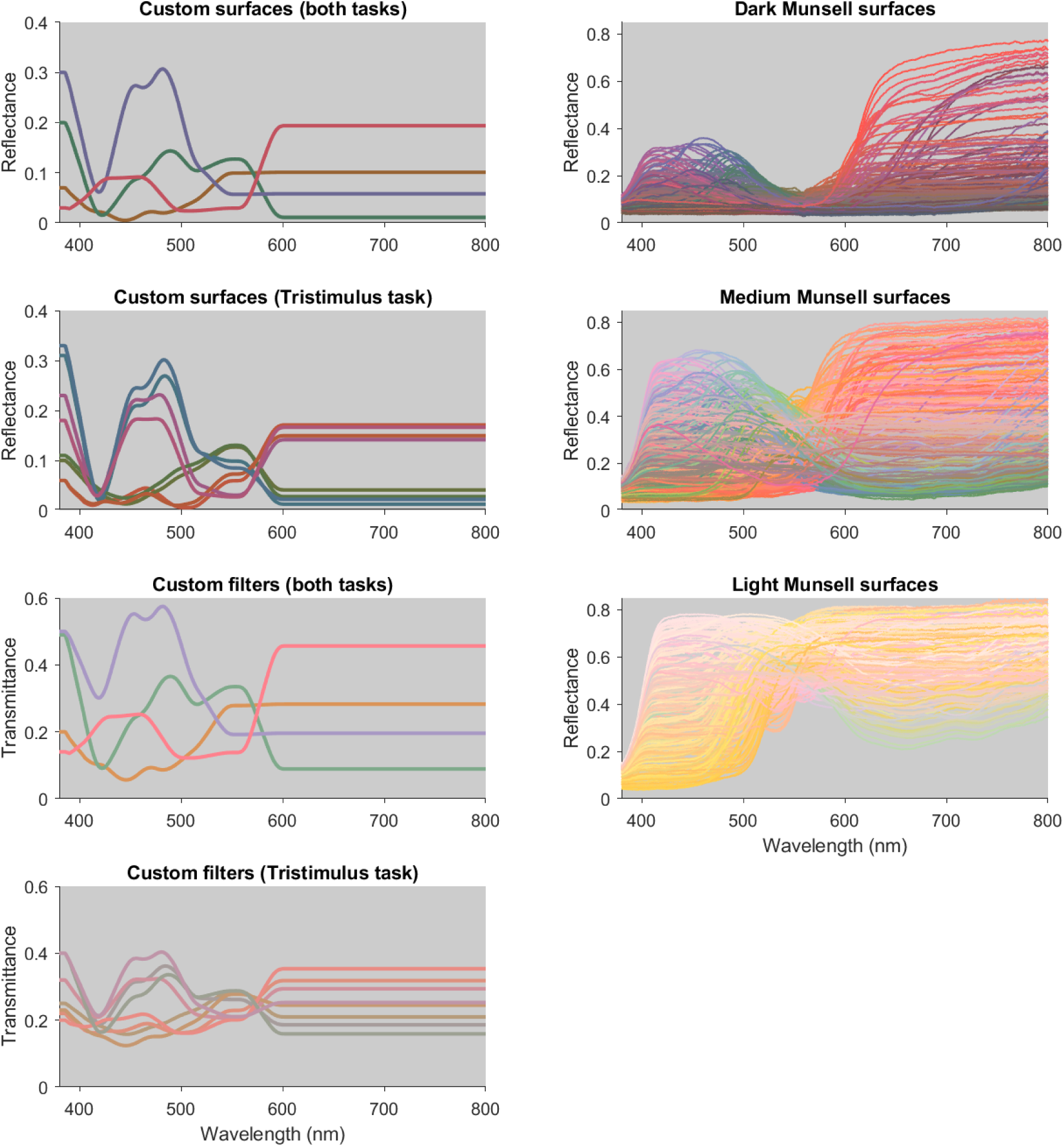
Surface reflectance functions and filter transmittance functions for all surfaces and filters used in the stimuli for the visual search task. In the left column, the custom surface reflectance and filter transmittance functions are shown, including all of those indicated in **Figure 1B-C**. There were 4 custom surfaces and 4 custom filters used in both the surface reflectance and tristimulus value search. To produce additional tristimulus matches, in the tristimulus value search there were a further 8 custom surface and 8 custom filters. In the right column, the Munsell surface reflectance functions are shown, divided into the three groups of values (dark, medium and light), that were assigned to all background surfaces. In each case, the functions are plotted in a colour approximating their appearance under a white illuminant (for surfaces) or when placed in front of white light (for filters).

## Notes

### Competing Interest Statement

The authors have declared no competing interest.

### Summary of Updates

Supplementary figure added, analysis from Exp 1 updated, new model added to modelling predictions

